# Root Hydraulic and Metabolic Regulation Drives Drought Tolerance in Napier Grass

**DOI:** 10.64898/2026.03.28.714958

**Authors:** Swee-Suak Ko, Yi-Chien Wu, Sy-Chyi Cheng, Min-Jeng Li, Tzu-Rung Li, Jeng-Bin Lin, Chi-Hui Sun, Charles C.-K. Chou, Kuo-Chen Yeh

**Author notes:** Corresponding author. Email address (K.C. Yeh) and (Charles C.-K. Chou). These authors contributed equally to this work (Swee-Suak Ko, Yi-Chien Wu, Sy-Chyi Cheng).

## Abstract

Napier grass (*Cenchrus purpureus* syn. *Pennisetum purpureum*), a perennial C_4_ forage and bioenergy crop, exhibits strong drought resilience, yet the integrative mechanisms underlying this tolerance remain incompletely understood. This study examined physiological, hydraulic, and metabolic responses of four Napier grass cultivars under PEG-induced osmotic stress and progressive soil water deficit. Drought significantly increased the root-to-shoot ratio, indicating preferential biomass allocation to roots, which supported maintenance of shoot growth and tissue water status. All cultivars showed an approximate twofold increase in water-use efficiency (WUE) under water deficit, with cv2 and cv7 displaying superior performance. Upregulation of *aquaporin* genes (*PIP2;2* and *PIP2;3*) suggested active hydraulic regulation that sustained carbon assimilation under reduced transpiration. Metabolic profiling revealed pronounced root-centered osmotic adjustment, including accumulation of galactinol, myo-inositol, raffinose family oligosaccharides, proline, and several amino acids. Enhanced expression of the *galactinol synthase* gene confirmed activation of raffinose biosynthesis pathways. Genotypic variation highlighted cv2 as particularly drought resilient. Rapid post-stress regrowth further underscored the importance of perennial root persistence. In conclusion, drought tolerance in Napier grass arises from coordinated hydraulic resilience, osmotic adjustment, and C_4_ photosynthetic efficiency, supporting its suitability for forage and bioenergy production in water-limited environments.

**Significant:** This study shows drought tolerance in Napier grass relies on root-driven hydraulic and metabolic regulation with efficient water-use efficiency, rather than avoidance, and that PEG responses predict field performance.

## Introduction

Drought is one of the most severe abiotic stresses limiting crop productivity worldwide, and its impact is expected to intensify under future climate scenarios characterized by higher temperatures and irregular precipitation patterns. C_4_ crops and grasses, including maize (*Zea mays*), sorghum (*Sorghum bicolor*), sugarcane (*Saccharum* spp.), and Napier grass (*Cenchrus purpureus*, syn. *Pennisetum purpureum*), exhibit higher intrinsic water-use efficiency than C_4_ plants due to their CO_2_-concentrating mechanism and reduced photorespiration. Nevertheless, drought remains a major constraint on biomass accumulation and yield stability in C_4_ species, particularly during early establishment and regrowth phases in perennial grasses (Ghannoum *et al*., 2010). Understanding the mechanistic basis of drought tolerance in C_4_ grasses is therefore critical for sustaining forage and bioenergy production under water-limited conditions.

Drought stress is commonly imposed experimentally as either water deficit or osmotic stress, yet these treatments represent distinct physiological challenges. Water deficit arises from progressive soil drying and imposes an integrated whole-plant stress that alters soil–root hydraulic conductance, reduces leaf water potential, promotes abscisic acid–mediated stomatal closure, and increases the risk of xylem cavitation. In contrast, osmotic stress is typically simulated using osmotica, such as polyethylene glycol (PEG), that lower external water potential without substantially altering oxygen diffusion. Although both treatments reduce cellular water availability, osmotic stress primarily induces rapid cellular dehydration responses and strong metabolic reprogramming, whereas water deficit more accurately reflects long-term developmental and hydraulic adjustments occurring under field conditions (Blum, 2010; Verslues *et al*., 2006). This distinction is particularly relevant for C_4_ grasses, in which sustained leaf hydraulic conductance and coordination between mesophyll and bundle sheath function are essential for maintaining photosynthesis during drought (Tardieu *et al*., 2018).

Regulation of plant water transport under drought relies heavily on aquaporins, a family of membrane proteins that facilitate transcellular water movement. In C_4_ crops, plasma membrane intrinsic proteins (PIPs) and tonoplast intrinsic proteins (TIPs) play key roles in controlling root hydraulic conductivity, leaf water status, and recovery after rehydration. Drought stress frequently induces isoform-specific regulation of aquaporins, with some PIPs being downregulated to restrict water loss, while others are maintained or induced to support cellular turgor and hydraulic continuity (Hachez *et al*., 2012; Maurel *et al*., 2015). In maize and sorghum, expression levels of *PIP1* and *PIP2* family members have been associated with drought tolerance and root water uptake efficiency, highlighting aquaporins as potential molecular markers for drought adaptation in C_4_ grasses (Abdel-Ghany *et al*., 2020; Vandeleur *et al*., 2009).

In parallel with hydraulic regulation, osmotic adjustment through the accumulation of compatible solutes constitutes a major drought tolerance strategy. Soluble sugars, including galactose-derived oligosaccharides, raffinose, and trehalose, contribute to osmotic balance, stabilization of membranes and proteins, and protection against oxidative damage. Raffinose family oligosaccharides accumulate in drought-tolerant grasses and have been shown to function as antioxidants and cellular protectants under dehydration (Nishizawa *et al*., 2008; Taji *et al*., 2002). Trehalose, although typically present at low concentrations, plays a regulatory role through trehalose-6-phosphate signaling, influencing carbon allocation and growth–stress trade-offs. Enhanced trehalose metabolism has been linked to improved drought tolerance and maintenance of photosynthetic capacity in C_4_ crops (Nuccio *et al*., 2015).

Amino acid metabolism also contributes significantly to drought tolerance, with proline and γ-aminobutyric acid (GABA) being particularly prominent. During drought, proline accumulation functions in osmotic adjustment, reactive oxygen species scavenging, and stabilization of proteins and membranes, and is frequently correlated with drought tolerance in maize and sorghum (Szabados and Savouré, 2010). GABA, produced via the GABA shunt, serves both metabolic and signaling roles, linking carbon–nitrogen balance with stress responses. Emerging evidence indicates that GABA modulates stomatal conductance, antioxidant capacity, and stress recovery, thereby contributing to drought tolerance in grasses (Bouche and Fromm, 2004). In addition to proline, PEG-induced osmotic stress promotes accumulation of glutamate-derived amino acids, asparagine, alanine, and branched-chain amino acids, collectively supporting osmotic adjustment, nitrogen remobilization, and metabolic flexibility in plants (Bowne *et al*., 2012).

Despite these advances, the mechanistic basis of drought tolerance in Napier grass, a high-biomass perennial C_4_ grass widely cultivated for forage and bioenergy, remains poorly defined. Most insights into C_4_ drought adaptation derive from annual cereals, whereas Napier grass exhibits distinct traits, including vigorous regrowth, perennial root systems, and strong biomass recovery after stress. In particular, the coordinated roles of sugars and amino acids in regulating dehydration tolerance and growth in Napier grass remain largely unexplored. Finally, most existing studies rely on short-term osmotic stress assays, which may not adequately capture the integrated hydraulic and developmental responses occurring under field-relevant water-deficit conditions (Verslues *et al*., 2006). Addressing these knowledge gaps is essential for identifying robust physiological and molecular markers of drought tolerance and for guiding breeding and biotechnological strategies to improve resilience and productivity of Napier grass under water-limited environments.

## Materials and methods

### Plant Materials

Four Napier grass cultivars/lines (cv2, cv4, cv7, and ALS5) were selected to evaluate genotypic variation in drought tolerance. The detailed agronomic characteristics of the four cultivars are summarized in **Table S1**. Among them, cv2 is the most widely cultivated forage Napier grass cultivar in Taiwan, while cv4 is characterized by a large stem diameter and high tiller fresh weight. cv7 is distinguished by its short internode length, which confers strong resistance to lodging. These three cultivars were developed and released by the Taiwan Livestock Research Institute (TLRI). ALS5 is a Napier grass germplasm line collected from the Ali Mountain region of Taiwan and maintained by Academia Sinica. ALS5 generally exhibits tall growth with relatively thin stems.

### PEG6000-Induced Osmotic Stress Experiment

A controlled osmotic stress experiment was conducted to evaluate the growth and physiological responses of Napier grass under polyethylene glycol (PEG6000)-induced water deficit conditions. Uniform, healthy Napier grass plants at the five-leaf stage were grown hydroponically in full-strength (1×) Kimura nutrient solution prior to stress imposition. Osmotic stress was imposed by supplying plants with the same nutrient solution supplemented with 10% (w/v) polyethylene glycol #6000 (PEG6000, XR-LPOL024, Sigma-Aldrich, USA), which corresponds to an estimated osmotic potential of approximately −0.5 MPa at 25°C. The PEG6000 solution was freshly prepared prior to application and thoroughly mixed to ensure a uniform and stable osmotic potential throughout the treatment period. Control plants received the same 1× Kimura nutrient solution without PEG addition. To prevent oxygen depletion in the root zone and avoid hypoxic stress, both control and PEG-treated nutrient solutions were continuously aerated using an air pump system during the experiment. For each genotype, six individual plants were grown as biological replicates. Plants were subjected to osmotic stress for a period of two weeks under a rain-proof plastic greenhouse at the Biotechnology Center in Southern Taiwan (AS-BCST) (23◦06′14.4′′ N 120◦17′31.2′′ E). All plants were monitored regularly for visible stress symptoms.

### Drought Stress Under Field

Plants were grown on October 28, 2024, in a rainproof plastic greenhouse at AS-BCST under ambient environmental conditions. For each cultivar/line, six individual plants were grown in the field as biological replicates. Spacing area per plant was 20 cm × 20 cm in sandy loam soil.

After establishment, plants were grown under well-watered conditions for seven weeks to ensure uniform vegetative development. At the start of the drought treatment, all plants were irrigated thoroughly and uniformly to field capacity. Irrigation was then completely withheld to induce a progressive soil water deficit over six weeks, until the soil water content at a depth of 10 cm decreased to approximately 10%. During the drought period, growth-related traits were measured to assess drought responses among genotypes. At the end of the drought treatment, shoots were harvested by cutting the top portion of each plant 10 cm above the soil surface. This standardized cutting height was used to simulate defoliation and to enable assessment of regrowth potential. Following harvest, drought-stressed plants were rewatered to field capacity to assess post-drought recovery and regrowth capacity. Rewatering was applied uniformly across all cultivars and lines, and plants were subsequently maintained under well-watered conditions for the remainder of the experiment. Recovery-related growth parameters were recorded seven weeks after rewatering to quantify regrowth performance and resilience following exposure to drought stress.

### Measurement of Plant Growth, SPAD Value, and Photosynthesis Parameters

Plant growth and physiological responses were evaluated at two time points: immediately before drought treatment (day 0), and after six weeks of drought stress. Plant height was measured from the soil surface to the tip of the tallest leaf to assess overall growth performance. Root length was measured to assess root system development under osmotic stress. Shoot and root fresh biomass were determined immediately after harvest to evaluate aboveground and below-ground growth responses, respectively. The root-to-shoot fresh weight ratio was calculated to evaluate changes in biomass allocation under osmotic stress. Wilting symptoms were visually assessed using a standardized wilting score, in which plants were scored based on leaf rolling, drooping, and loss of turgor, with higher scores indicating more severe stress. Plant water content was determined by measuring fresh weight immediately after harvest and dry weight after oven-drying samples at 70°C until constant weight. Water content was calculated as the percentage of water relative to fresh weight. Biomass allocation patterns were assessed by calculating the root-to-shoot fresh weight ratio.

To further assess potential structural changes in leaves induced by osmotic stress, leaf mass per area (LMA) was measured. LMA is widely used as an indicator of leaf thickness, tissue density, and overall leaf structural investment in response to environmental conditions. For each sample, fully expanded leaves (L1) were collected, and their surface area was determined using image analysis. LMA was calculated using the following formula: LMA=Leaf mass/Leaf area. Each treatment consisted of six biological replicates (n = 6), and all measurements were conducted according to standardized protocols to ensure consistency and reliability across samples.

Leaf chlorophyll content was estimated using a SPAD chlorophyll meter. Measurements were taken from the L1, and three readings per leaf were recorded and averaged to obtain a representative SPAD value for each plant. Leaf gas exchange parameters were measured using a portable photosynthesis system (LI-6800F Gas Exchange and Fluorescence System; LI-COR Biosciences, Lincoln, NE, USA), following the procedure described by (Chen *et al*., 2022). Measurements were conducted on fully expanded, healthy leaves under steady-state conditions. The LI-6800 chamber was set to a leaf temperature of 30°C, a photosynthetic photon flux density of 1500 µmol mL² sL¹, an ambient CO_2_ concentration of 400 ppm, and a relative humidity of 60%. Net CO_2_ assimilation rate (A), stomatal conductance to water vapor (gsw), and transpiration rate (E) were recorded after instrument stabilization. All measurements were performed within the same time window each day to minimize diurnal variation. Instantaneous water-use efficiency (WUE) was calculated as the ratio of net CO_2_ assimilation rate to transpiration rate (A/E). All measurements were performed on 6 independent plants per treatment, which served as biological replicates (n = 6).

### Analysis of sugars and amino acids in Napier grass using ultra-performance liquid chromatography–tandem mass spectrometry (UPLC-MS/MS)

#### Chemicals and sample preparation

Acetonitrile (ACN, LC grade), ammonium hydroxide (NH_4_OH), methanol (MeOH), and hydrochloric acid (HCl) were purchased from J.T. Baker (Phillipsburg, NJ, USA). Amino acids and γ-aminobutyric acid (GABA) were obtained from Sigma–Aldrich (St Louis, MO, USA). Formic acid was obtained from Fluka (Germany). Deionized water was produced by a Milli-Q system (Millipore, Bedford, MA, USA). The amino acid and GABA standards were dissolved in 1 M HCl aqueous solution and MeOH, respectively, and then further diluted with MeOH to prepare calibration solutions at different concentrations (0.5–500 ppb).

Four Napier grass genotypes were grown under controlled environmental conditions and subjected to PEG-induced osmotic stress for two weeks. Root and fully expanded leaf tissues were harvested separately, immediately frozen in liquid nitrogen, and stored at −80°C until analysis. Each treatment consisted of three independent biological replicates per genotype (n = 3). The samples were ground to a fine powder in liquid nitrogen and then extracted with a 50% acetonitrile aqueous solution (50 mg/mL). After sonication in an ultrasonic bath for 1 h and centrifugation at 18,800×g at 4°C for 10 min, the supernatants were filtered through 0.2 μm polytetrafluoroethylene (PTFE) membrane filters for subsequent sugar and amino acid analysis.

#### Analysis of sugars in Napier grass

The ultra-performance liquid chromatography–high resolution tandem mass spectrometry (UPLC–HRMS/MS) system used for sugar analysis was as described previously (Sun *et al*., 2023). Briefly, 5 μL of extract solution was injected into a BEH Amide column (130Å, 1.7 µm, 2.1 × 100 mm, Waters, USA) installed on an UltiMate 3000 UHPLC system (Thermo Fisher Scientific, San Jose, USA). The column temperature was maintained at 35L. The mobile phases consisted of 80% ACN containing 0.1% NHLOH (eluent A) and 30% ACN containing 0.1% NHLOH (eluent B). The gradient started at 0% B, increased to 40% B over 10 min, ramped to 60% B over 3 min, held for 7 min, and then returned to 0% B within 1 min. For recycling, the initial gradient composition (0% B) was restored and maintained for 14 min. The flow rate was set at 0.15 mL/min during sample separation and increased to 0.3 mL/min during the recycling step.

Negative ions generated by a heated electrospray ionization (HESI) source were analyzed using an Orbitrap Fusion Lumos Tribrid mass spectrometer (Thermo Fisher Scientific) to acquire mass spectra at both MS1 (full scan, *m/z* 100–750) and MS2 (data-dependent acquisition, DDA) levels. The MS1 and MS2 scan resolutions were set at 60,000 and 30,000, respectively; and the parameters of the HESI source were as follows: ESI voltage, - 2.8 kV; sheath gas, 35 arbitrary units (Arb); auxiliary gas, 7 Arb; sweep gas, 1 Arb; ion transfer tube temperature, 300; and vaporizer temperature, 275. Raw data were processed using Compound Discoverer 3.3 software (Thermo Fisher Scientific), in combination with an online database (mzCloud) and an in-house sugar database (MS2 spectra and retention times), to identify sugars in the extracts and determine their peak areas.

#### Analysis of amino acids in Napier grass

An ExionLC UPLC system (Sciex, MA, USA) coupled with a triple quadrupole mass analyzer (QTRAP 6500+, Sciex) was utilized to quantify amino acids in the extracts. The injection volume was set at one μL, and an ACQUITY Premier BEH C18 VanGuard FIT column (Waters, 1.7 µm, 2.1 × 150 mm) was maintained at 40°C for chromatographic separation. The mobile phase comprised water with 0.1% formic acid (eluent A) and ACN with 0.1% formic acid (eluent B). The LC gradient started at 0% B for 1 min, increased to 10% B over 4.5 min, then ramped to 99% B over 4 min and held for 4.3 min, followed by a decrease to 0% B over 0.2 min. For recycling, the initial gradient composition (0% B) was restored and maintained for 4 min. The flow rate was set at 0.25 mL/min during sample separation and increased to 0.3 mL/min during the re-equilibration step. The eluent from LC system was delivered into an IonDrive Turbo V source for ionization and MS detection. The ion source settings were as follows: ionspray voltage, +5500LV; temperature (TEM), 500°C; curtain gas, 30L(arb); ion source gas 1 (GS1), 40 (arb); ion source gas 2 (GS2), 50 (arb). The mass spectrometer was operated in multiple reaction monitoring (MRM) mode (**Table S2**) to characterize fragment ions. Data processing was carried out using Sciex OS 3.0.0.3339 software.

### Real-time PCR

To investigate the effects of polyethylene glycol (PEG)–induced osmotic stress on gene expression in Napier grass, the transcript levels of selected stress-responsive marker genes were analyzed using quantitative real-time PCR (RT-qPCR). Total RNA was first reverse-transcribed into complementary DNA (cDNA), which was subsequently used as the template for real-time PCR amplification. The analyzed genes included those involved in proline and other amino acids biosynthesis, sugar metabolism, and water transport aquaporin genes (*PIP2;2, PIP2;3*). Gene expression was compared between roots and leaves to assess tissue-specific transcriptional responses to osmotic stress.

Total RNA was extracted from samples using TRIzol reagent (Promega, USA) according to the manufacturer’s instructions. First-strand cDNA was synthesized from 1 μg of total RNA using an M-MLV reverse transcription kit (Promega, USA) with oligo(dT) primers, following standard protocols. The reverse transcription reaction was performed under the recommended conditions, and the resulting cDNA was diluted 20× for subsequent quantitative PCR analysis. Quantitative

PCR (qPCR) assays were conducted using gene-specific primers (listed in **Table S3**) under optimized amplification conditions. Each reaction was carried out in a total volume of 15 μL, containing 7.5 μL of SYBR Green master mix (KAPA Biosystems, Merck), forward and reverse primers, and a cDNA template. Amplification was performed using a CFX Duet Real-Time PCR System (Bio-Rad, USA). Relative gene expression levels were calculated using the 2 ΔΔCt method. Ct values of target genes were normalized against the geometric mean of two internal reference genes, *18S rRNA* and *Actin*, to obtain ΔCt values, ensuring stable normalization across samples. ΔΔCt values were then calculated relative to the control condition. To evaluate treatment effects, gene expression levels were further reported as the ratio of PEG-treated to control (CK) samples (PEG/CK ratio), providing a comparative measure of transcriptional responses under stress conditions. All reactions were performed in triplicate (biological and technical) to ensure reproducibility and accuracy.

### Statistical Analysis

Statistical significance was determined using one-way analysis of variance (ANOVA), followed by Duncan’s multiple range test (DMRT) at *p* ≤ 0.05. Means sharing the same letter are not significantly different, whereas means with different letters indicate significant differences. Student’s *t*-test was used to compare differences between control (CK) and drought-treated plants.

## RESULTS

### PEG-induced osmotic stress reveals intrinsic drought-responsive traits in Napier grass

To identify intrinsic drought-responsive mechanisms independent of field heterogeneity, four Napier grass genotypes (cv2, cv4, cv7, and ALS5) were evaluated under 10% PEG-induced osmotic stress. Two weeks of PEG6000 treatment (−0.5 MPa) significantly affected plant growth and biomass allocation, revealing clear genotypic differences. Controls plants (CK) exhibited normal growth, whereas PEG-treated plants showed visible wilting symptoms and growth inhibition. Among the genotypes, cv2 exhibited the least growth retardation and developed longer roots under osmotic stress (**Fig. 1**). Plant height was not significantly different between CK and PEG-treated cv2, whereas the other three genotypes showed a significant reduction in plant height (**Fig. 2A**). In contrast to shoot growth, root length increased significantly in cv2 following PEG treatment (P < 0.05; **Fig. 2B**). PEG treatment reduced shoot biomass across all genotypes except cv2 (**Fig. 2C**). In contrast, root biomass was less affected, with cv2 and cv4 maintaining higher root biomass than cv7 and ALS5 under both control and PEG-treated conditions (**Fig. 2D**). As a result, the root-to-shoot ratio increased significantly under osmotic stress, particularly in cv4 and cv7 (**Fig. 2E**), indicating a shift in biomass allocation toward root development. After two weeks of PEG treatment, all genotypes exhibited a significant increase in leaf mass per area (LMA), suggesting increased leaf thickness (**Fig. 2F**). In addition, PEG treatment led to a significant increase in wilting scores (**Fig. 2G**) and a reduction in leaf water content in all genotypes, although water content remained above 73% (**Fig. 2H**).

**Fig. 1.**
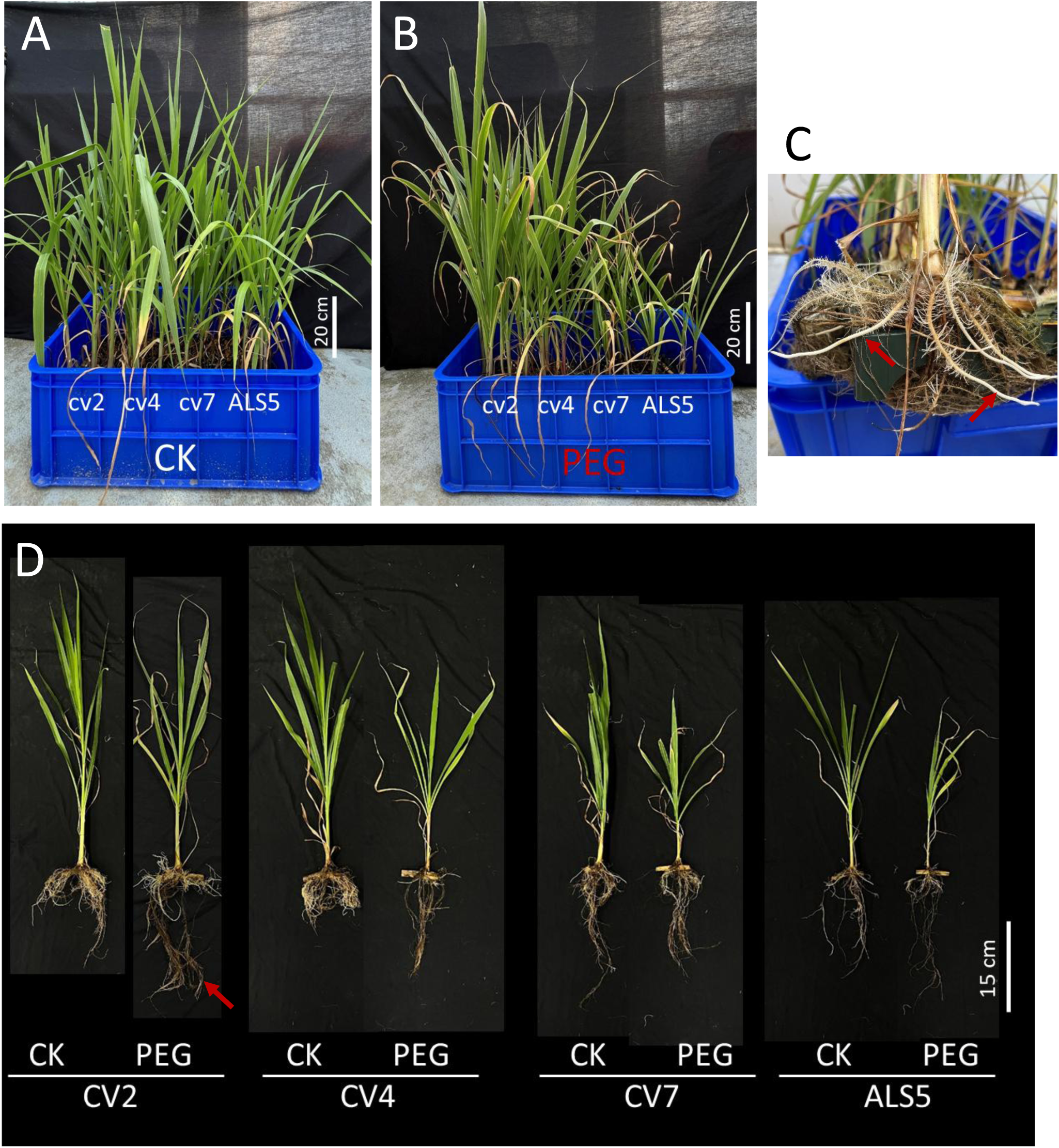
PEG-induced osmotic stress responses in four Napier grass cultivars. (A) Control plants grown without PEG treatment in 1× Kimura nutrient solution. (B) Plants exposed to 10% (w/v) PEG 6000 for two weeks, exhibiting visible symptoms of osmotic stress. (C) Evidence of PEG-induced osmotic adjustment in cultivar cv2, characterized by the emergence of new roots under stress conditions (arrows). (D) Comparative phenotypic analysis of the four Napier grass cultivars after two weeks of PEG treatment versus their respective controls grown in 1× Kimura nutrient solution.

**Fig. 2.**
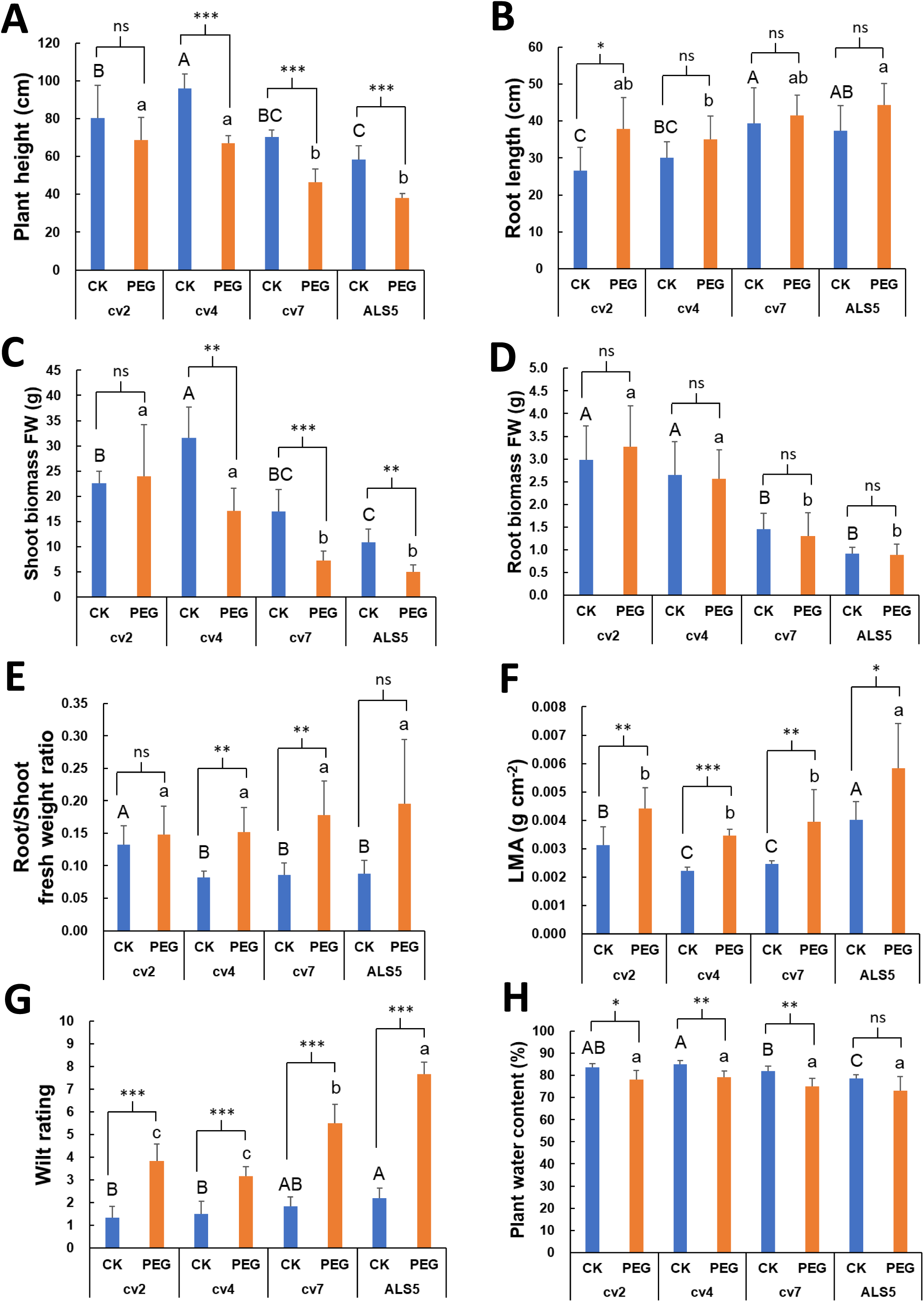
Growth responses of Napier grass subjected to 10% PEG6000-induced osmotic stress for 2 weeks. (A) Plant height measured after osmotic stress. (B) Root length, indicating root system development under osmotic stress. (C) Shoot fresh biomass weight, representing aboveground growth and water status. (D) Root fresh biomass, showing below-ground growth changes. (E) Root-to-shoot fresh weight ratio, illustrating biomass allocation shifts under stress. (F) Leaf mass per area (LMA), indicating changes in leaf thickness or density. (G) Wilting score, assessing the severity of stress symptoms visually. (H) Plant water content, measuring hydration status post-treatment. Sample size: n = 6 biological replicates. Different uppercase letters indicate significant differences among genotypes under control conditions (CK), as determined by Duncan’s multiple range test (DMRT; p < 0.05). Different lowercase letters indicate significant differences among genotypes under PEG stress, as determined by DMRT (p < 0.05). Student’s t-test was used to assess significant differences between the CK and PEG treatments. Asterisks indicate significance levels: * p < 0.05, ** p < 0.01, *** p < 0.001, and n.s. indicate no significant difference. Error bars represent the standard error of the mean (n = 6).

Leaf chlorophyll content, as estimated by SPAD values, significantly decreased under PEG stress in all genotypes; however, cv2 and cv4 maintained higher SPAD values compared to the other genotypes, indicating delayed chlorophyll degradation under osmotic stress (**Fig. 3A**). Aside from cv2, all genotypes showed significant reductions in CO_2_ uptake, stomatal conductance (gsw), transpiration rate (E), and electron transfer rate (ETR) (**Fig. 3B–E**) after PEG stress. Interestingly, cv2 and cv7 maintained similar water-use efficiency (WUE) after PEG stress, while cv4 and ALS5 exhibited a significant decrease (**Fig. 3F**).

**Fig. 3.**
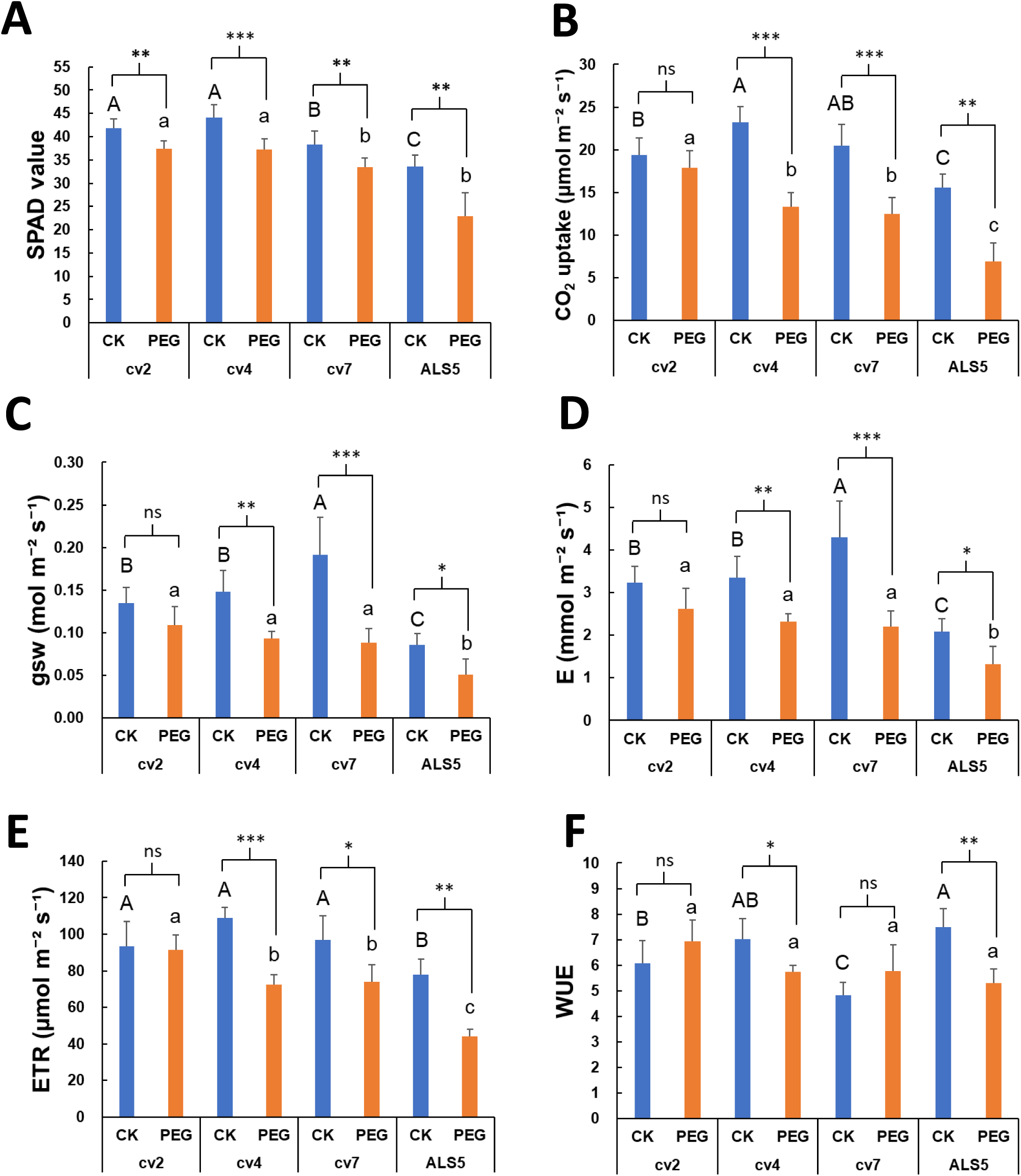
Chlorophyll and photosynthesis of Napier grass subjected to 10% PEG6000-induced osmotic stress. (A) Chlorophyll content measured using a SAPD meter. (B) CO_2_ uptake. (C) Stomatal conductance (gsw). (D) Transpiration rate (E). (E) Electron transfer rate (ETR). (F) Water-use efficiency (WUE). Plant tissues were harvested two weeks after treatment with 10% polyethylene glycol (PEG). Sample size: n = 6 biological replicates.

### Sugar metabolism in Napier grass under PEG-induced osmotic-stress

Sugar composition profiles were analyzed separately in leaf and root tissues to resolve tissue-specific metabolic responses to PEG-induced osmotic stress. UPLC–HRMS/MS analysis revealed substantial remodeling of sugar metabolism following PEG treatment, with pronounced, and in some cases contrasting, responses between leaf and root tissues as well as among genotypes (**Fig. S1A, B**). In roots, osmotic stress induced a marked accumulation of galactose-derived sugars, particularly galactinol and myo-inositol. The cv7 and ALS5 accumulated trehalose in roots, but it was reduced in their corresponding leaf tissues (**Fig. S1A,B**). Trisaccharides and tetrasaccharides accumulated to higher levels in roots than in leaf tissues (**Fig. S1C, D**), indicating differential regulation of sugar metabolism between organs.

Notably, sucrose content in roots decreased tremendously in cv2 and cv7 under PEG treatment, whereas myo-inositol and galactinol levels increased in the roots of cv2, cv4, and ALS5. In contrast, trehalose accumulation declined in cv2 but increased in cv7 and ALS5 (**Fig. 4**), highlighting both shared and genotype-specific root metabolic adjustments to osmotic stress. Consistent with these metabolic changes, gene expression analysis revealed PEG-stress transcriptional responses in roots. The *sucrose-phosphate synthase* gene (*SPS3*) was downregulated, suggesting reduced sucrose biosynthesis; while the *galactinol synthase* (*GolS2*) and *myo-inositol monophosphatase1* (*MIP1*) genes, key genes in galactinol and myo-inositol biosynthesis, were significantly upregulated. Additionally, expression of the *trehalose-6-phosphate synthase* gene (*TPS1*) was slightly elevated in cv7 roots (**Fig. 4G**). Together, these transcriptional adjustments correlated with the observed accumulation of osmoprotective sugars, indicating a coordinated root-specific response to osmotic stress.

**Fig. 4.**
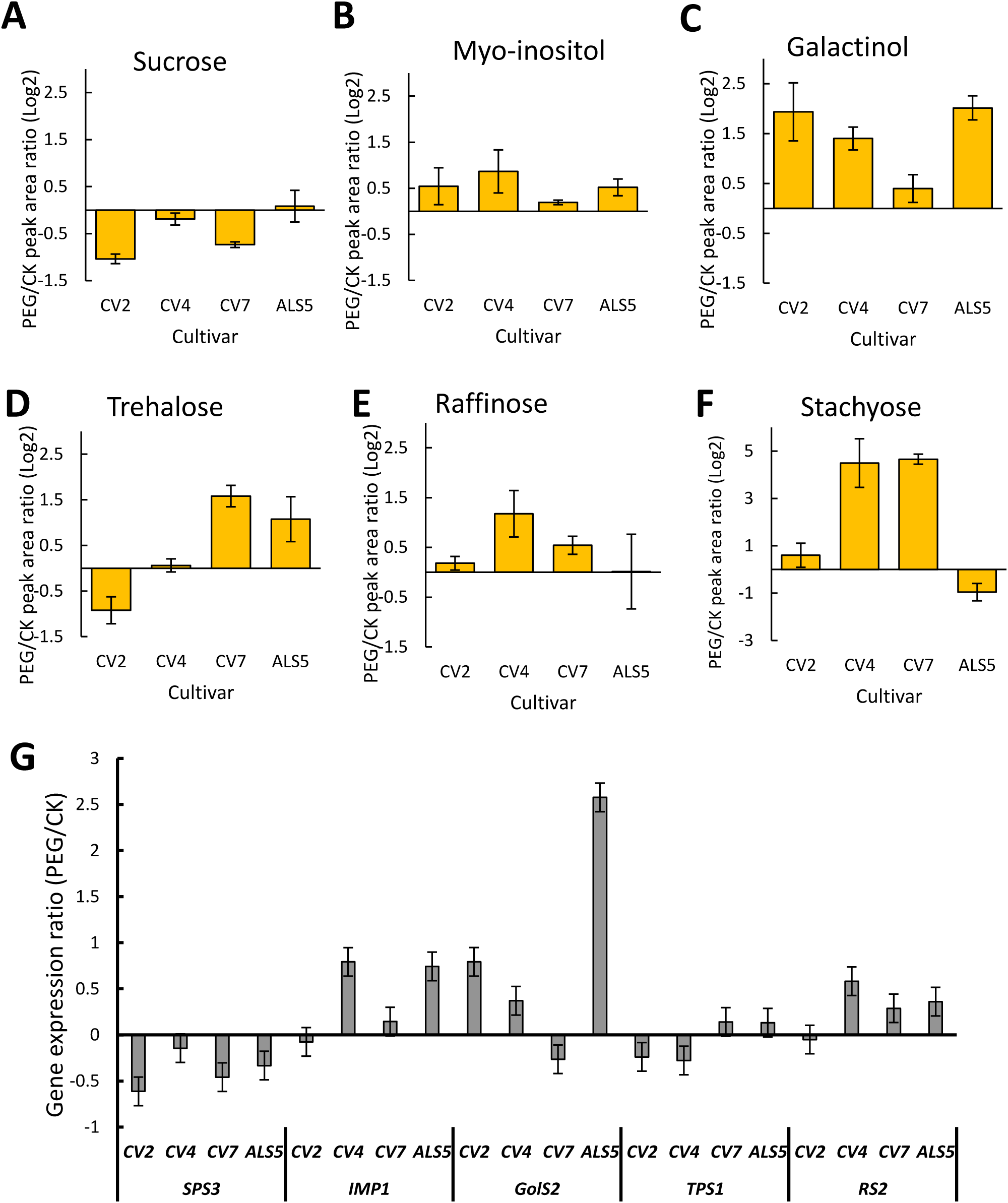
Changes in sugars and sugar biosynthesis gene expression in the roots of four Napier grass genotypes under PEG-induced drought stress. Ultra-performance liquid chromatography–high-resolution tandem mass spectrometry analysis was used to characterize soluble sugars, including: (A) sucrose, (B) myo-inositol, (C) galactinol, (D) trehalose, (E) raffinose, and (F) stachyose following 10% PEG6000 treatment for two weeks. Data are presented as logL fold changes (PEG/CK), calculated as the ratio of PEG-treated samples to controls (CK). (G) Expression patterns of genes involved in sugar biosynthesis, presented as PEG-to-CK ratio. *SPS3*, *sucrose-phosphate synthase3*; *MIP1, myo-inositol monophosphatase1*; *GolS2*, *galactinol synthase2*; *TPS1*, *trehalose-6-phosphate synthase1*; and *RS2*, *raffinose synthase2*. Values represent the means ± SD of three independent biological replicates (n = 3).

### Amino acid metabolism in Napier grass under PEG-induced osmotic stress

UPLC–MS/MS was used to profile amino acid metabolites in leaf and root tissues under PEG-induced osmotic stress and control conditions. The analysis revealed pronounced, tissue-specific changes in amino acid metabolism in response to PEG treatment. Overall, the majority of amino acids showed greater accumulation in root tissues than in leaves under osmotic stress, indicating a stronger metabolic response in roots (**Fig. S2**). Notably, asparagine (Asp) and glutamine (Gln) were remained relatively unchanged in leaves (**Fig. S2A**), whereas their levels significantly decreased in roots following PEG treatment (**Fig. S2B**). In roots, cv2 showed the highest accumulation of histidine, proline, methionine, isoleucine, and leucine under PEG-induced osmotic stress, whereas ALS5 exhibited the lowest levels of these amino acids relative to the other cultivars (**Fig. 5**). In addition, GABA levels were significantly elevated in roots under PEG-induced osmotic stress, highlighting a pronounced root-specific metabolic response to stress conditions (**Fig. S3**).

**Fig. 5.**
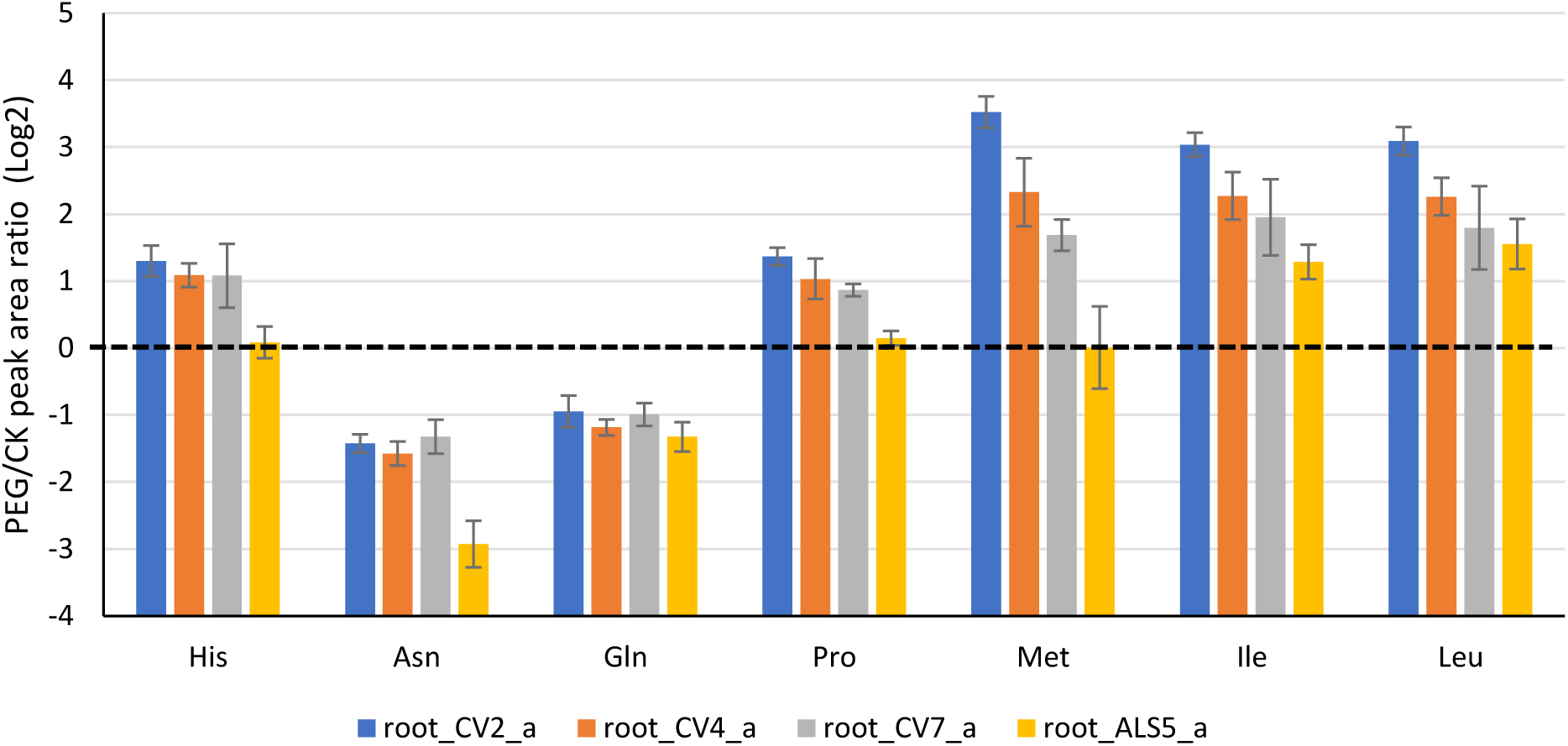
Polyethylene glycol (PEG) treatment altered the composition of amino acid metabolites in roots under PEG-induced stress. Root tissues were harvested two weeks after treatment with 10% PEG. Values represent the means ± SD of three independent biological replicates (n = 3).

### PEG-induced osmotic stress upregulates aquaporin gene expression

The expression of two *plasma membrane intrinsic protein* genes, *PIP2;2* and *PIP2;3*, was examined in leaf and root tissues under PEG-induced osmotic stress. Gene expression analysis revealed that *PIP2;2* was upregulated in both roots and leaves of most genotypes, except in the root of cv2 and the leaves of ALS5 (**Fig. 6A**). In contrast, *PIP2;3* expression was significantly upregulated in both roots and leaves of all tested genotypes, with the exception of the leaves of ALS5. Notably, *PIP2;3* expression in the roots of cv7 increased approximately fivefold under PEG treatment compared with the control (CK) (**Fig. 6B**).

**Fig. 6.**
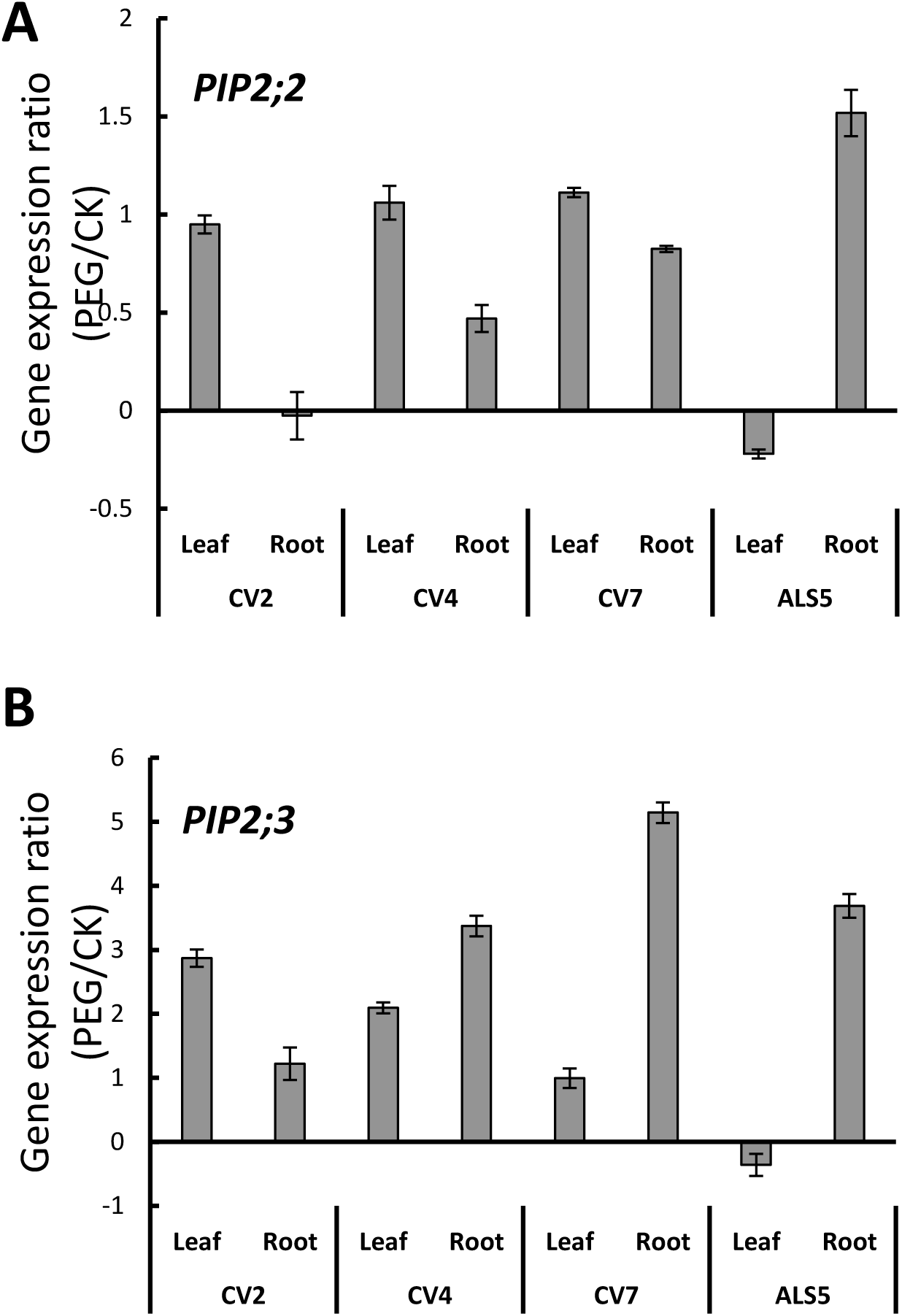
Gene expression patterns of *aquaporin* genes in the roots and leaves of four Napier grass genotypes in response to polyethylene glycol (PEG)-induced osmotic stress. Relative expression levels under PEG treatment compared with the control (CK) are shown for (A) *PIP2;2* and (B) *PIP2;3*. Data represent mean ± standard error from three independent biological replicates (n = 3).

### Genotypic divergence in biomass production and water-use efficiency under field water deficit

To validate PEG-defined drought-responsive traits under agronomically relevant conditions, the same genotypes (cv2, cv4, cv7, and ALS5) were evaluated under progressive soil water deficit in the field (**Fig.7**). After establishment under well-watered conditions, irrigation was withheld for six weeks, leading to a marked reduction in soil water content and visible wilting symptoms in all genotypes. However, cv2 and cv4 exhibited less severe wilting than cv7 and ALS5 (**Fig.7B**).

**Fig. 7.**
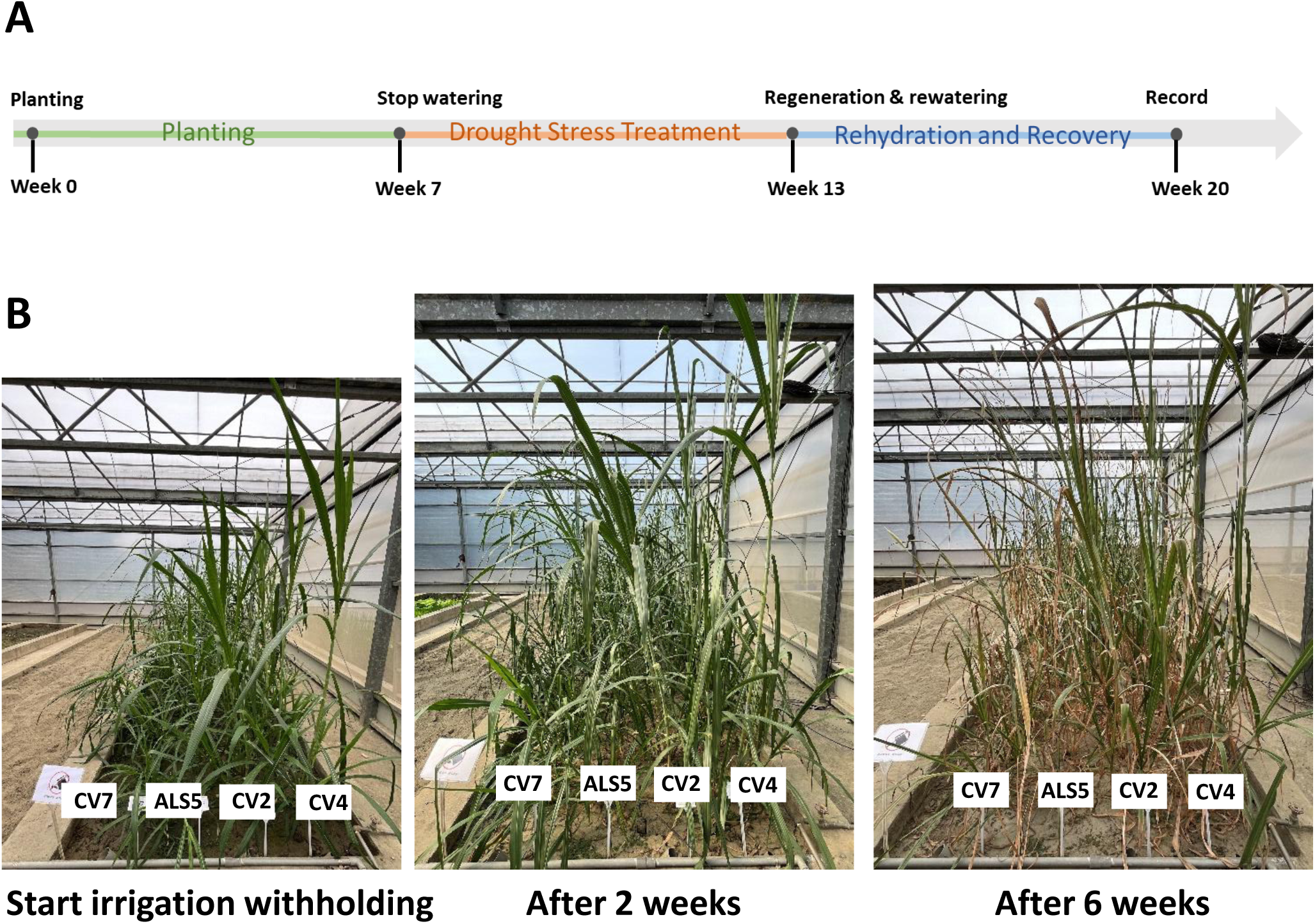
Experimental setup and phenotypic response of plants subjected to drought treatment. (A) Illustration of the drought treatment protocol applied to the plants, showing the timing and conditions of water deprivation. (B) Phenotypic differences between control plants (well-watered) and drought-stressed plants after the drought treatment period, including visible signs of wilting, leaf curling, and overall plant vigor reduction.

Photosynthesis measurements revealed genotype-specific physiological responses to drought. Prior to stress imposition, all four cultivars exhibited similar CO_2_ assimilation rates, indicating comparable photosynthetic capacity under well-watered conditions. However, carbon assimilation was significantly reduced under reduced soil moisture, declining to approximately half of pre-stress levels across all cultivars (**Fig. 8A**). Stomatal conductance (gLw) and transpiration rates decreased markedly in response to drought, with mean values reduced to approximately one-quarter of pre-stress levels after six weeks (**Fig. 8B, C**). Despite these reductions, intrinsic water-use efficiency (WUE), calculated as the ratio of CO_2_ assimilation to transpiration, increased under drought conditions. The most pronounced increases (approximately twofold) were observed in cv2 and cv4, whereas ALS5 showed a smaller response (**Fig. 8D**). Moreover, ΦPSII, electron transport rate (ETR), and non-photochemical quenching (NPQ) were significantly decreased during drought stress (**Fig. 8E-G**).

**Fig. 8.**
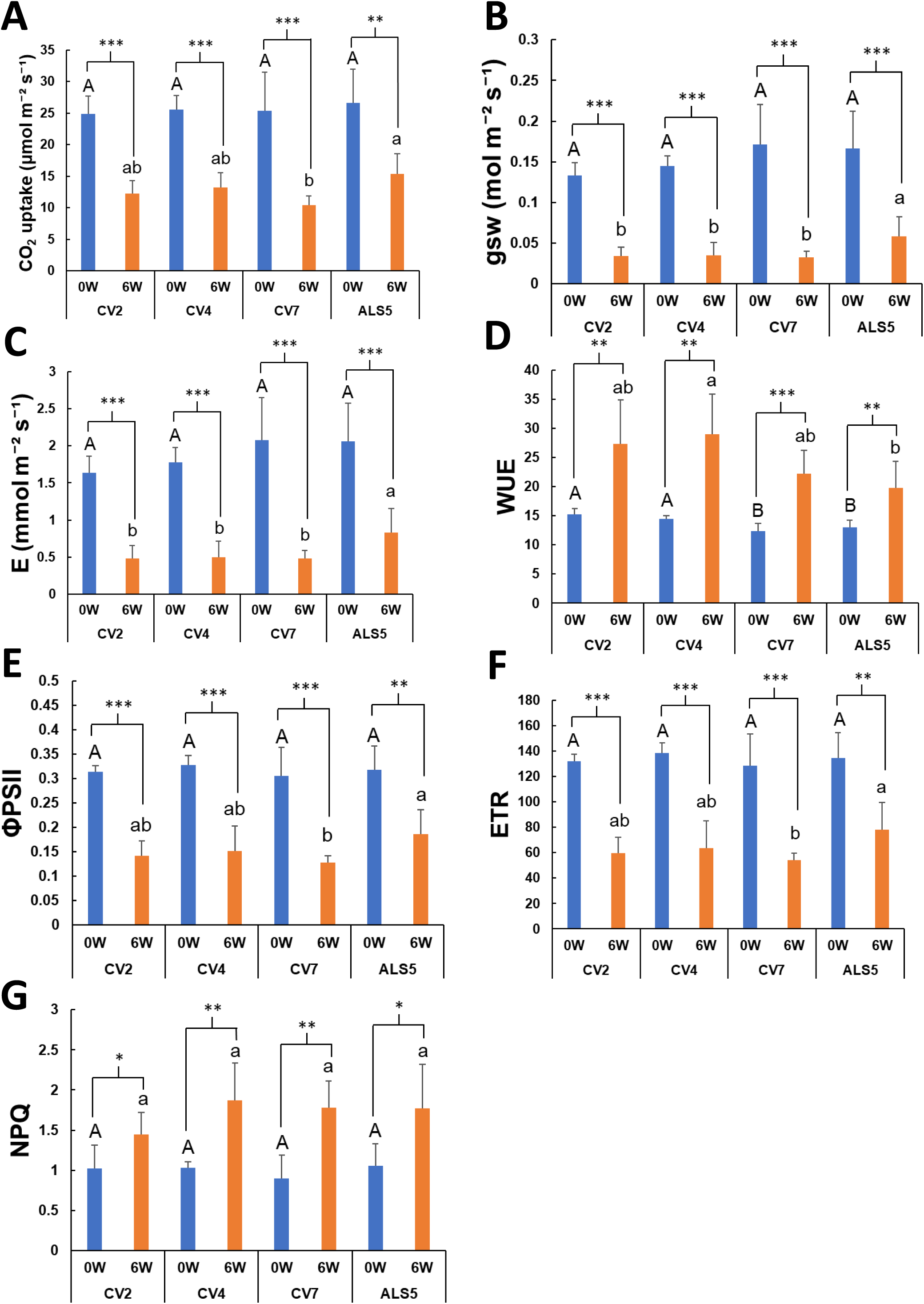
Photosynthesis parameters on the first day and after drought treatment for six weeks. (A) CO_2_ uptake. (B) Stomatal conductance (g_s_w). (C) Transpiration rate (E). (D) Water-use efficiency (WUE) calculated by A/E. (E) ΦPSII. (F) Electron Transport Rate (ETR). (G) Non-photochemical quenching (NPQ). Statistical significance was determined by a one-way analysis of variance (ANOVA), followed by Duncan’s multiple range test (DMRT). Values followed by a different letter(s) are significantly different at p < 0.05. Error bars indicate the SD of the mean from six plants.

**Table 1** compares the growth performance of four Napier grass genotypes before and after six weeks of irrigation withholding. Prior to stress, cv4 and ALS5 exhibited the greatest plant height, followed by cv2, whereas cv7 showed the shortest stature. Under drought conditions, ALS5 remained the tallest, followed by cv4 and cv2, while cv7 remained the shortest (**Table 1**), indicating genotypic variation in drought-induced growth inhibition. Leaf chlorophyll content, estimated using SPAD values, also differed significantly among genotypes. Cv2 consistently exhibited the highest SPAD values both before and after drought stress, suggesting superior chlorophyll retention and delayed leaf senescence under water-limited conditions. Interestingly, cv7 showed an increase in SPAD values following drought treatment and exhibited the highest tiller number after stress. Overall, biomass accumulation was significantly affected by both drought stress and genotype. Cv4 produced the highest fresh and dry biomass under drought conditions, followed by cv2 and ALS5, whereas cv7 accumulated the lowest biomass. The reduced biomass of cv7 was largely associated with its shorter plant height and lower shoot mass. Plant water content after six weeks of drought differed significantly among genotypes. Cv2 and cv4 maintained significantly higher plant water content (77%) than cv7 and ALS5 (74%), indicating superior water retention capacity under prolonged soil drying (**Table 1**)

**Table 1.** Effects of withholding irrigation for six weeks on the agronomic traits of Napier grass.

| Variety | Plant Ht. (cm) |  | SPAD value |  | Tiller No. | Biomass FWt | Biomass DWt | Water content |
| --- | --- | --- | --- | --- | --- | --- | --- | --- |
|  | Before | After | Before | After |  | (g/ plant) | (g/ plant) | (%) |
| **CV2** | 95B | 125c | 61A | 57a | 3.8b | 329b | 74b | 77.5a |
| **CV4** | 116A | 163b | 57B | 54ab | 4.2b | 473a | 107a | 77.6a |
| **CV7** | 38C | 75d | 35C | 50b | 5.7a | 119d | 30d | 73.7b |
| **ALS5** | 107AB | 195a | 58B | 43c | 3.8b | 195c | 52c | 74.6b |

Statistical significance was determined using one-way analysis of variance (ANOVA), followed by Duncan's multiple range test (DMRT). Values followed by different letters indicate significant differences at $p < 0.05$ . Uppercase letters denote differences before drought treatment, while lowercase letters indicate differences after six weeks of withheld irrigation. Error bars represent the standard deviation (SD) of the mean from six plants.

### Regrowth capacity of Napier grass following drought and cutting

As a perennial species with a well-developed root system, Napier grass exhibited strong regrowth capacity following cutting and rewatering across all cultivars. Plants were cut to a uniform height of 10 cm above the soil surface at week six, after which regrowth was monitored over time. New shoots emerged within 10 days in all genotypes, and fully expanded leaves were observed at 15 and 34 days after cutting (**Fig. 9**). After seven weeks of regrowth, all genotypes exhibited similarly high SPAD values, indicating comparable leaf chlorophyll content. However, significant genotypic differences were observed in shoot traits. Cv2 plants were the tallest, whereas cv7 and ALS5 were the shortest. Shoot dry biomass also varied significantly: cv4 produced the highest biomass, followed by cv2 and cv7, whereas ALS5 accumulated the least. Tiller number during regrowth varied significantly among cultivars. Cv7 produced the highest number of tillers, averaging 26.7 tillers per plant, whereas cv2, cv4, and ALS5 produced fewer than 15 tillers per plant (**Table 2**).

**Fig. 9.**
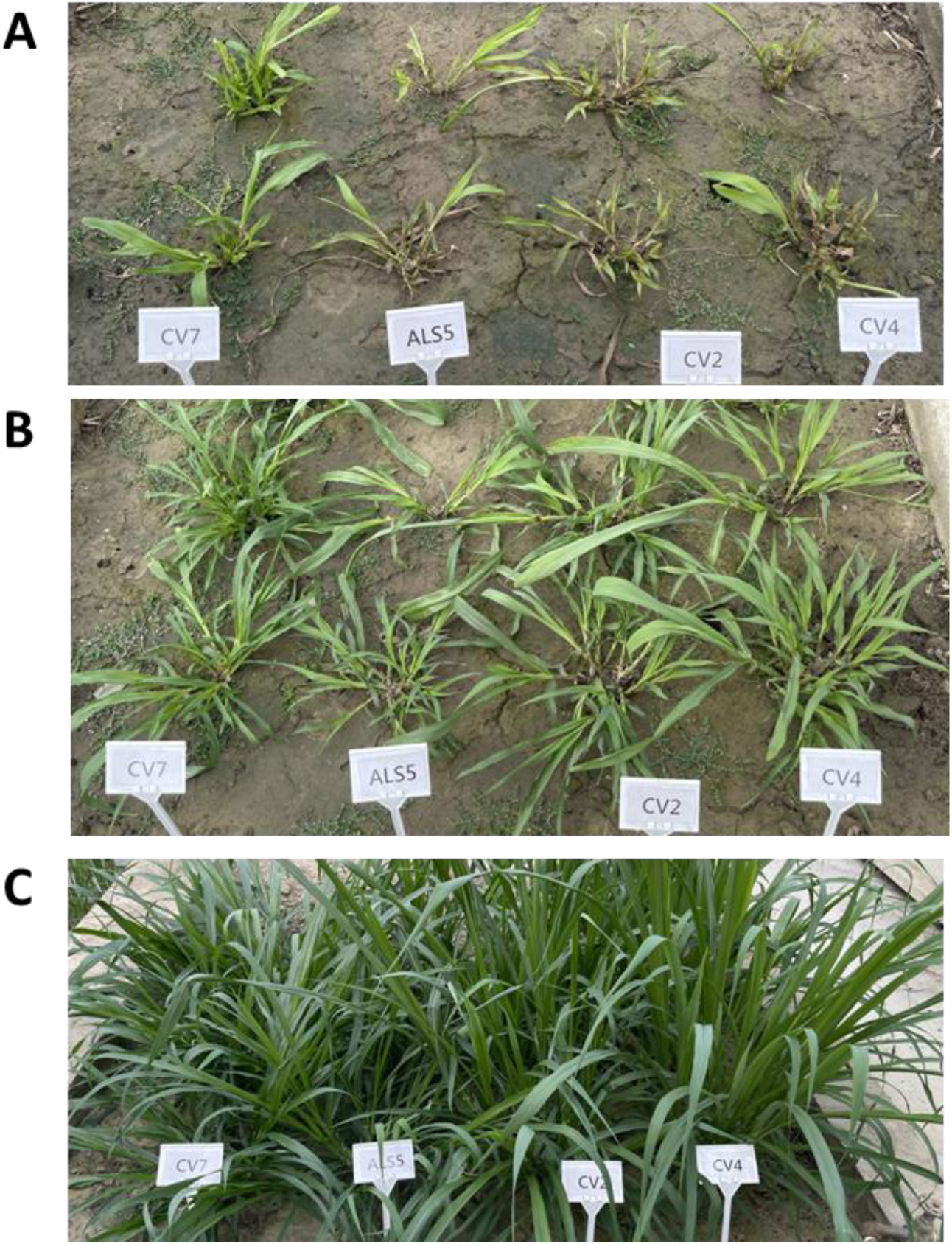
Regrowth performance of Napier grass following a period of drought and subsequent recovery irrigation. (A) Plant regrowth 10 days after resuming irrigation, demonstrating initial recovery. (B) Plant regrowth status 15 days post-recovery irrigation, showing continued improvement in biomass and vigor. (C) Regrowth of Napier grass 34 days after irrigation restoration, illustrating near-full recovery or sustained growth following drought stress.

**Table 2.**
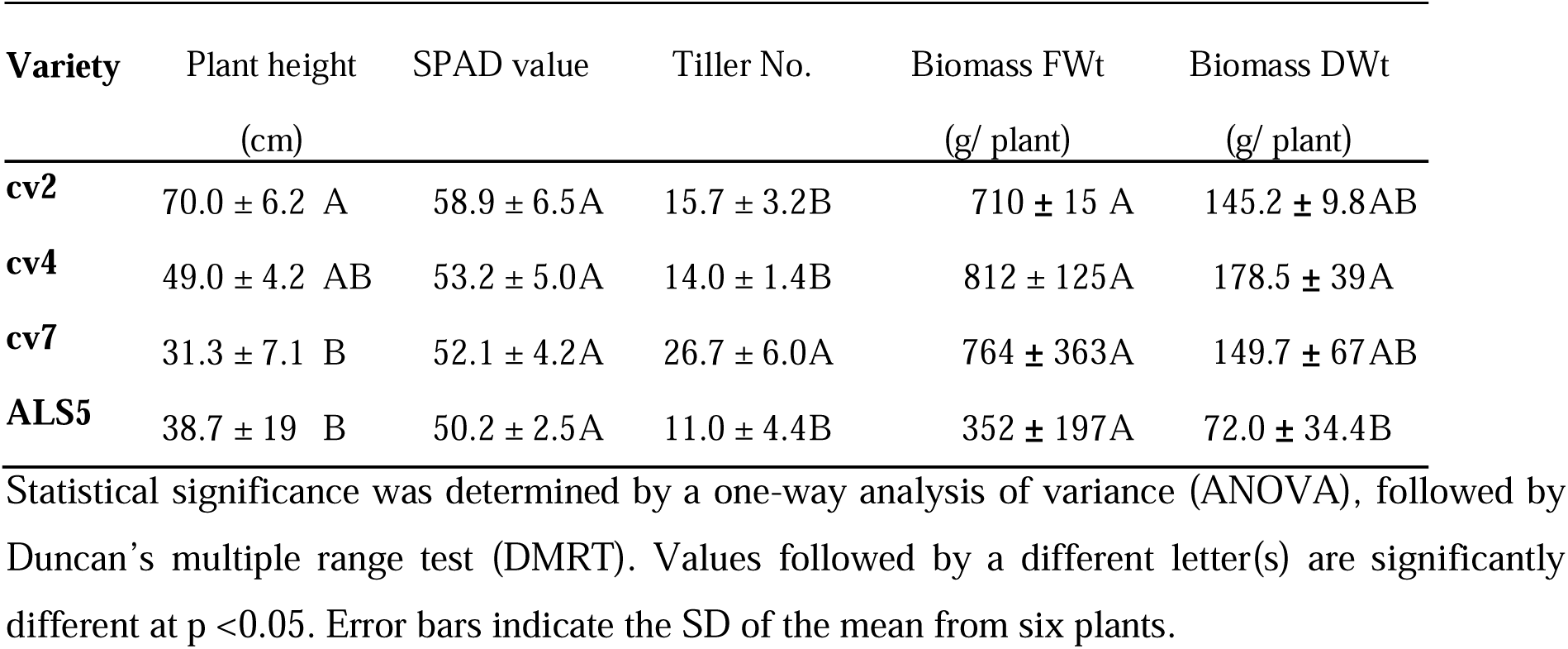
Effects of seven-week recovery following the withholding of water on agronomic traits of Napier grass.

| **Variety** | Plant height<br>(cm) | SPAD value | Tiller No. | Biomass FWt<br>(g/ plant) | Biomass DWt<br>(g/ plant) |
| --- | --- | --- | --- | --- | --- |
| **cv2** | 70.0 ± 6.2 A | 58.9 ± 6.5A | 15.7 ± 3.2B | 710 ± 15 A | 145.2 ± 9.8AB |
| **cv4** | 49.0 ± 4.2 AB | 53.2 ± 5.0A | 14.0 ± 1.4B | 812 ± 125A | 178.5 ± 39A |
| **cv7** | 31.3 ± 7.1 B | 52.1 ± 4.2A | 26.7 ± 6.0A | 764 ± 363A | 149.7 ± 67AB |
| **ALS5** | 38.7 ± 19 B | 50.2 ± 2.5A | 11.0 ± 4.4B | 352 ± 197A | 72.0 ± 34.4B |

Statistical significance was determined by a one-way analysis of variance (ANOVA), followed by Duncan's multiple range test (DMRT). Values followed by a different letter(s) are significantly different at p <0.05. Error bars indicate the SD of the mean from six plants.

## Discussion

C perennial Napier grass exhibits a suite of drought-adaptive strategies that distinguish it from annual C_4_ cereals and C_4_ crops. These traits reflect its perennial growth habit, high biomass demand, and adaptation to water-limited environments. In this study, PEG-induced osmotic stress significantly increased the root-to-shoot ratio (**Fig. 2E**), indicating strategic biomass reallocation toward below-ground tissues. Such shifts enhance water acquisition and are a well-established adaptive response to drought (Lynch, 2013). Because shoot growth is typically more sensitive to declining water potential than root elongation, preferential investment in roots enables access to deeper or more stable soil moisture. This reduces whole-plant hydraulic resistance, buffers decline in leaf water potential, and helps sustain stomatal conductance and photosynthesis under stress (Vadez, 2014). Consistently, enhanced root development in Napier grass was associated with better maintenance of shoot growth and higher tissue water content during drought (**Fig. 2E, H**).

Water-use efficiency (WUE) further distinguished drought responses among genotypes. Under PEG-induced stress, cv2 and cv7 maintained higher WUE than the other cultivars (**Fig. 3F**), and all four cultivars exhibited an approximate twofold increase in WUE under progressive soil water deficit (**Fig. 8D**). Increased WUE under drought generally reflects coordinated adjustments in stomatal conductance and carbon assimilation (Chaves *et al*., 2003). Intrinsically, C_4_ species possess 1.5–3 times higher WUE than C_4_ crops due to their CO_2_-concentrating mechanism and reduced photorespiration (Ghannoum *et al*., 2010; Sage and Zhu, 2011). While annual C_4_ crops such as maize are highly productive under adequate water, their shorter life cycles and limited post-drought recovery constrain resilience under prolonged stress (Campos *et al*., 2004). In contrast, the perennial growth of Napier grass supports sustained water uptake, enhanced buffering capacity, and rapid recovery following stress, conferring advantages in drought-prone environments. Similarly, other perennial C_4_ species, such as Miscanthus, exhibit strong drought tolerance (Al Hassan *et al*., 2022).

This performance is underpinned by the C_4_ photosynthetic pathway, which enables high CO assimilation at lower stomatal conductance. Napier grass utilizes the NADP-ME subtype (Neto and Guerra, 2019), which may provide a strong electron sink and maintain photochemical efficiency as stomatal conductance declines, thereby sustaining carbon gain during water stress. The integration of C_4_ photosynthesis coupled with Napier grass’s robust root system explains its superior drought performance relative to C_4_ crops such as rice and wheat.

Hydraulic regulation is central to the resilience of Napier grass. Aquaporins, particularly plasma membrane intrinsic proteins (PIPs), regulate root hydraulic conductivity and enable rapid adjustments in water transport under stress (Maurel *et al*., 2015). Drought stress upregulated *PIP2;2* and *PIP2;3* in Napier grass roots and leaves (**Fig. 6**), indicating active hydraulic regulation rather than a passive consequence of increased root biomass. By reducing internal hydraulic resistance, aquaporins help sustain carbon assimilation at lower transpirational cost, further enhancing intrinsic WUE (Hachez *et al*., 2012). This coordination of root hydraulics and C_4_ photosynthesis allows Napier grass to maintain functional photosynthesis while limiting water loss.

Osmotic adjustment constitutes another key drought-adaptive mechanism. Under reduced water potential, plants maintain turgor and metabolic activity through the accumulation of compatible solutes, particularly soluble sugars and amino acids (Blum, 2010). In Napier grass, this response was predominantly root-centered. PEG-induced stress led to greater accumulation of soluble sugars and amino acids in roots than in leaves, with significant increases in galactinol and myo-inositol (**Fig. 4B, C**). Galactinol is a precursor of raffinose family oligosaccharides (RFOs), which stabilize membranes and proteins during stress, while myo-inositol contributes to osmotic regulation, antioxidant defense, and signaling (Valluru and Van den Ende, 2011). Preferential root accumulation of these metabolites likely supports cellular homeostasis and root function, thereby sustaining shoot performance. Consistent with this, RFO biosynthesis was activated under drought. RFOs—including raffinose, stachyose, and verbascose—are synthesized from sucrose using galactinol as a galactosyl donor, with galactinol synthase (GolS) acting as a rate-limiting enzyme. *GolS* expression is widely induced by abiotic stress and correlates with stress tolerance (Nishizawa *et al*., 2008; Taji *et al*., 2002). In this study, *GolS* transcripts were upregulated in roots under PEG stress (**Fig. 4G**), confirming activation of the RFO pathway. Metabolic profiling further showed more pronounced drought-induced reprogramming in roots than in leaves (**Fig. S1, S2**), reinforcing the root as the primary site of stress adaptation. Compatible solutes such as proline, GABA, raffinose, and galactinol accumulated in roots, contributing to osmotic balance, membrane protection, and redox homeostasis (Nishizawa *et al*., 2008; Szabados and Savouré, 2010).

Beyond proline, several amino acids—including histidine, methionine, isoleucine, and leucine—increased in Napier grass roots under water deficit (**Fig. 5**). Accumulation of specific amino acids during drought is associated with osmotic adjustment, redox buffering, and metabolic regulation (Handa *et al*., 1986; Kavi Kishor *et al*., 2015). Branched-chain amino acids (BCAAs), including isoleucine and leucine, are recognized as important components of stress-induced metabolic reprogramming, functioning not only as alternative respiratory substrates but also as signaling molecules under carbon–nitrogen imbalance (Less and Galili, 2008; Urano *et al*., 2009). Under drought conditions, when photosynthetic carbon assimilation is constrained, enhanced BCAA accumulation may support mitochondrial respiration by feeding into the tricarboxylic acid (TCA) cycle, thereby sustaining ATP production and cellular energy status. In parallel, BCAAs and their catabolic intermediates may act as metabolic signals that coordinate stress-responsive gene expression and nitrogen redistribution. Methionine metabolism provides an additional regulatory layer, as it serves as a precursor for ethylene and polyamines—both central regulators of stress perception and response (Joshi *et al*., 2010). Increased methionine availability may therefore promote ethylene-mediated transcriptional reprogramming and polyamine-dependent stabilization of membranes and macromolecules, enhancing cellular tolerance to dehydration. Histidine accumulation further contributes to stress adaptation by improving antioxidant capacity, potentially through metal ion chelation and activation of reactive oxygen species (ROS)-scavenging systems, thereby mitigating oxidative damage under drought stress (Ji *et al*., 2022; Khan *et al*., 2019). These metabolic adjustments appear to be transcriptionally coordinated, which likely drives the observed accumulation of these metabolites in Napier grass. Such integration of transcriptional regulation with metabolic flux suggests an active reallocation of nitrogen resources toward protective and energy-sustaining pathways. This coordinated response is consistent with drought-induced metabolic shifts reported in other grasses (Obata *et al*., 2015), and underscores a conserved mechanism in which amino acid metabolism supports both osmotic adjustment and energy homeostasis. Collectively, these processes likely enhance the capacity of Napier grass to maintain cellular function and recover following water deficit, contributing to its overall drought resilience.

Genotypic variation further shaped drought responses. Cv2—the most widely cultivated cultivar in Taiwan—showed less shoot inhibition, enhanced root elongation, and greater accumulation of protective metabolites such as galactinol and amino acids (**Figs. 4C, 5**). This combination of physiological and metabolic traits underscores substantial plasticity within Napier grass and highlights opportunities for selection and improvement. A defining feature of Napier grass is its rapid post-stress recovery. The perennial root system supports strong regrowth after rewatering or cutting, with new shoots emerging within 10 days and fully expanded leaves within 15–34 days (**Fig. 9**). This contrasts with many annual C_4_ crops, which often exhibit prolonged growth suppression after drought. Sustained root function, osmotic adjustment, and metabolic stability during stress collectively enable this rapid restoration of photosynthetically active tissue (Blum, 2010; Tardieu *et al*., 2018).

Taken together, drought tolerance in Napier grass arises from coordinated integration of three strategies: (1) hydraulic resilience supported by a plastic root system and aquaporin-mediated water transport; (2) root-centered osmotic adjustment through accumulation of compatible solutes, including RFOs and amino acids; and (3) strong regrowth capacity enabled by perennial root persistence. Rather than relying solely on drought avoidance, Napier grass maintains carbon assimilation, preserves leaf water status, and rapidly resumes growth following stress. These traits constitute valuable physiological targets that can be harnessed to improve drought resilience in perennial C_4_ forage and bioenergy crops.

In conclusion, Napier grass combines root-dominated hydraulic and metabolic adjustments with C_4_ photosynthetic efficiency and rapid regrowth to thrive under water-limited conditions. Its integrative drought tolerance framework highlights its value as both a resilient forage crop and a model for understanding stress adaptation in high-biomass perennial C_4_ systems.

## Data availability

Data supporting the findings of this study are available within this paper and supplementary data.

## Acknowledgments

We gratefully acknowledge the funding from the Academia Sinica Alpha project—Application of Agricultural Biomass for Carbon Sink Generation and Green Energy (AS-KPQ-112-NETZ-06-A) that supports this study. We thank the Taiwan Livestock Research Institute for providing Napier grass cultivars for this work. We also thank the genetically modified greenhouse of the Biotechnology Center of Southern Taiwan (AS-BCST) for providing the greenhouse and technical support of Yueh-Hua Lin; and mass spectrometry core facility in AS-BCST for their technical assistance. We are grateful to Ms. Miranda Loney for assistance with English editing.

## Supplementary data

**Table S1.** Agronomic traits of Napier grass used in this study.

**Table S2.** Multiple reaction monitoring (MRM) transitions and parameters for the detection of amino acids.

**Table S3.** Primers used in this study.

**Fig. S1.** Sugar and sugar alcohol composition in leaves and roots of four Napier grass genotypes under PEG-induced drought stress.

**Fig. S2.** PEG-induced changes in amino acid metabolism in Napier grass.

**Fig. S3.** PEG-induced changes in γ-aminobutyric acid (GABA) in the root and leaf of four Napier grass cultivars.

